# Network centrality is the most important variable determining gene sequence evolution rates in *Schizosaccharomyces pombe*

**DOI:** 10.1101/2021.01.10.426089

**Authors:** Simon Harnqvist

## Abstract

Which variables determine the rate of gene and protein sequence evolution is a central question in evolutionary genomics. In the model organism fission yeast *(Schizosaccharomyces pombe)*, the determinants of the rate of sequence evolution have yet to be determined. Previous studies in other organisms have typically found gene expression levels to be most significant, with numerous other variables identified as having a smaller impact. Here, partial least squares regression (PLS) and partial correlation analysis are used to model sequence evolution rates in the fission yeast genome by a range of variables. Variable importance in projection (VIP) scores as well as partial correlation coefficients are calculated for each variable, and used as estimates of the influence of each independent variable on sequence evolution rate. Unlike many previous studies in other organisms, centrality in the PPI network is shown to be the most important variable, and gene expression found to be less influential. Considerable heterogeneity is found in the influence of different gene ontology terms as well as amino acid composition. However, the majority of variance in constraint in fission yeast remains unexplained by this study, indicating that variables not yet considered and stochastics probably have considerable impact on the rate of molecular evolution.

## BACKGROUND

The question of which variables determine the rate of sequence evolution in both genes and proteins is one of the most central in evolutionary genomics. While there is a long list of variables that are believed to influence the rate of sequence evolution, the importance of each has still not been explored in fission yeast. The importance of each variable has been quantitatively examined – notably in humans by Alvarez-Ponce and colleagues (2017) – using regression techniques to estimate the influence of each variable. Because of the importance of fission yeast (*Schizosaccharomyces pombe)* in genetic research, it is important to investigate whether the drivers of molecular evolution differ in fission yeast compared to other organisms.

Kimura and Ohta (1974) suggested, based on the neutral theory of molecular evolution (Kimura 1968) that functional importance (importance of gene for organismal fitness) would be the most important predictor of sequence of evolution constraint. This hypothesis is intuitive; mutations in highly important genes should be more detrimental to fitness, providing a selective pressure against sequence change. However, once confounders are adjusted for, it seems that functional importance only has a minor impact on sequence evolution rates (Zhang and Yang 2015). Instead, gene expression level has arisen as the seemingly most important factor determining sequence conservation – this is true overall in most organisms (Zhang and Yang 2015) and a strong correlation has indeed been shown in fission yeast (Mata 2003). There are at least four hypotheses to why this is. The protein misfolding avoidance hypothesis (Drummond et al. 2005) proposes that translational robustness (and therefore slower evolution) results from a need to avoid toxic misfolding, the consequences of which are worse the more molecules of the protein are in a cell. The protein misinteraction avoidance hypothesis (Yang et al. 2012) suggest that, similarly, the cost of random misinteraction increases with protein concentration, meaning that the selective pressure to prevent such interactions increases with expression. The mRNA folding requirement hypothesis (Park et al. 2013), suggests that mutations in highly expressed mRNAs would have worse consequences for the strongly folded mRNAs of highly expressed genes compared to the more weakly folded mRNAs of less expressed genes. Finally, the expression cost hypothesis (Cherry 2010; Gout et al. 2010), proposes that because a mutation reducing a protein’s fitness by a fraction *k* is equivalent to a loss of the same fraction *k* protein molecules, the fitness loss to the organism would be greater with highly expressed proteins. It is likely that several of these mechanisms contribute at least to some extent to the correlation between constraint and expression, making it apparent why this correlation is so strong, relatively speaking.

Many other predictors of conservation have been identified. These include centrality in the protein-protein interactions (PPI) network (Ingram 1961; Fraser and Hirsh 2004; Jovelin and Phillips 2009; Alvarez-Ponce et al. 2017), although one study that compared the budding yeast, worm, and fly genomes (Hahn and Kern 2005) failed to detect any such correlation. The relationship reported in most studies is that proteins that have more interaction partners, or are more central by other metrics, evolve more slowly. This, as first suggested by Ingram (1961), is generally thought to be because mutations in highly connected proteins could disrupt more pathways than in less connected proteins. Additionally, a host of variables influencing the rate of sequence evolution in a range of organisms has been reported before, including but not limited to protein length (Lipman et al. 2002), pleiotropy (Hahn and Kern 2005), intron presence (Carmel et al. 2007), cellular location (Liao et al. 2010), and codon bias (Drummond and Wilke 2008); Zhang and Yang (2015) provide a comprehensive review of the various variables implicated in sequence evolution.

Determining how much of the variance in sequence evolution rates is determined by each variable is a challenging high-dimensional regression problem. Previous studies (e.g. Jovelin and Phillips 2009; Yang and Gaut 2011; Alvarez-Ponce et al. 2017) have done this, often using principal component regression (PCR); however, the models produced by the studies cited explain less than 20% of the variance in sequence evolution rates. Instead, the present study proposes the use of partial least squares regression (PLS), a method that maximises the variance in the dependent variable (y) that is explained by the independent variables (X) (Haenlein and Kaplan 2004). Finding the determinants of the rate of molecular evolution involves determining which variables explain how much variance in that rate, making PLS a suitable choice. This technique offers a method of easily quantifying the influence of each variable in the regression by calculating the variable importance in projection (VIP) (Mehmood et al. 2012). Additionally, partial correlation analysis, which finds the correlation between two variables adjusting for the influence of covariates, is used as a second estimate of the influence of each variable.

The present study uses publicly available sequence evolution and functional genomics data from Grech and colleagues (Grech et al. 2019), with additional data from the PomBase (Lock et al. 2019) and STRING (Szklarczyk et al. 2019) databases, to construct a PLS model of the rate of sequence evolution in the fission yeast genome. The VIP scores are then extracted and used as a metric of how important each variable is as a predictor of sequence evolution constraint. At time of writing, there appears to be no readily available data on protein sequence evolution in *S. pombe*; this article does nonetheless discuss sequence evolution in both protein and DNA sequences.

## METHODS

### Sequence evolution constraint calculations

Grech and colleagues (2019) aligned the genome of *Schizosaccharomyces pombe* with those of three other fission yeasts (*S. japonicus, S. octosporus, and S. cryophilus*), and used the phyloP algorithm (Siepel et al 2006, cited in Grech et al. 2019) to estimate sequence evolution constraint per gene.

### Data sources

The gene list (n = 5609) of most protein coding genes in the fission yeast genome, with data on constraint (calculated as above), gene and protein expression expression, gene length, chromosome, essentiality and solid media fitness, was retrieved from Grech and colleagues (2019), retrieved from their article’s Figshare repository. Interactome data were retrieved from STRING. Gene ontology data were retrieved using the GO Term Mapper (go.princeton.edu/cgi-bin/GOTermMapper), with the gene list from Grech et al. as input. All other data, including amino acid composition and protein length and size, were retrieved from PomBase (Lock et al. 2019). The links to the datasets are available under “Original data sources”

### Data pre-processing

Gene ontology (GO) annotation terms as well as chromosome data were one-hot encoded using the mltools R package (Gorman 2018); all other variables were already numeric. Amino acid composition data were scaled to proportion of protein. Missing data were imputed using the missForest R package (Stekhoven and Buhlmann 2012). All preprocessing was carried out in R 4.0.0 (R Core Team 2020) with extensive use of the Tidyverse packages (Wickham et al. 2019).

### Network analysis

Betweenness centrality, closeness centrality, degree centrality (all three proposed by Freeman 1979), and eigenvector centrality (Bonacich 1972), were calculated for each gene based on protein interactions data with the igraph (Csardi and Nepusz 2006) R package, using a minimum STRING interaction score of 0.400. Betweenness centrality measures how often a node is on the shortest path in a network, closeness centrality how far the node is from all other nodes, degree centrality how many direct neighbours a node has, whereas eigenvector centrality measures how many other important nodes a node is connected to.

### Partial least squares regression modelling

Partial least squares regression (PLS/PLSR) was chosen to model constraint by all other variables because it is suitable for multivariate non-parametric modelling with noisy and non-orthogonal variables (Cramer 1993; Mehmood et al. 2012). It also allows easy interpretation of variable influence in the model by calculating the variable importance in projection (VIP) score for each variable (Mehmood et al. 2012). The model was trained with the pls R package (Mevik et al. 2019) using all variables except gene name and constraint as independent variables (X), with constraint as the dependent variable (y) to be predicted. All data were centred and scaled. Ten-fold cross-validation was used in training to guard against overfitting, and a 20% random holdout set was used to verify that performance on non-training data was comparable to that on training data. The optimum number of components was chosen with the selectNcomp function in the pls package.

### Variable importance in projection

Variable importance in projection scores were calculated using the PLSVarSel package (Mehmood et al. 2012). VIP can informally be described as the influence each independent variable has on each component of the model, adjusting for the variance in the independent variable explained by that component; the mathematical detail is well covered by Chong and Jun (2005). A rule of thumb is that variables with a VIP of less than 1 can be excluded from a model (Chong and Jun 2005) – however, the purpose here is not variable selection, but importance inference.

### Model comparisons

As comparison with the PLS model, principal component regression (PCR) and random forest (RF) models were trained. The PCR model was trained with the pls package (Mevik et al. 2019), whereas the Random Forest model was trained using randomForest (Liaw and Wiener 2002). The number of components to keep in the PCR model was selected with selectNcomp. The random forest model was trained using the default hyperparameters of 500 trees and mtry of number of variables divided by 3. Root mean squared error on the 20% holdout test set was calculated for each model using the rmse function in the Metrics package (Hamner and Frasco 2018). Variance explained by PLS/PCR models was calculated with the R2 function in pls. Variance explained by the random forest model was calculated using ‘pseudo R^2^’ (Liaw and Wiener 2002):

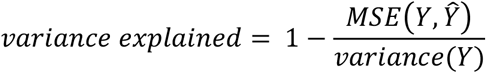

Where MSE is the mean squared error of prediction, Y is the independent variable vector, and Ŷ the vector of predicted values.

### Partial correlation analysis

Partial correlations were calculated between each independent variable and constraint (phyloP score), using all other variable groups as covariates. Note that other variables within the same group were not used as covariates because this would decrease the apparent correlation where there is collinearity that we do not wish to adjust for (e.g. degree and closeness centrality; adjusting for one removes the correlation of the other). Kendall’s rank correlation coefficient was chosen as a nonparametric alternative to Pearson’s correlation coefficient. Calculations were done using the Pingouin package (Vallat 2018) in Python 3.8.5.

## RESULTS AND DISCUSSION

### Network centrality is the most important determinant of constraint, followed by gene expression, functional importance, and sequence length

With three of the highest variable importance in projection scores (fig. 1a) as well as the two strongest partial correlations with sequence evolution constraint (fig. 1b), network centrality measures emerge as the strongest predictors of constraint. Closeness centrality shows the strongest correlation with a partial correlation coefficient (τ) of 0.12, whereas degree centrality displays the highest VIP score (3.4). While this result is not entirely unexpected, and in line with relatively recent findings in humans (Alvarez-Ponce et al. 2017), this does contrast against the consensus that gene expression is the most dominant variable (Zhang and Yang 2015). One likely explanation this is the improvement in quality and quantity in interactions data in the past decade, allowing better inferences. It is possible that there is no biological discrepancy between fission yeast and other examined organisms with regards to whether network centrality influences constraint. If anything, one might speculate that the importance of conserving protein-protein interactions might be even *higher* in multicellular organisms, where pathways between cells are more interconnected, as opposed to in yeasts. However, the results show that gene expression is still a highly important variable, with a relatively high VIP score of 2.2 and a partial correlation coefficient (τ) of 0.09 with constraint.

**Figure 1.**
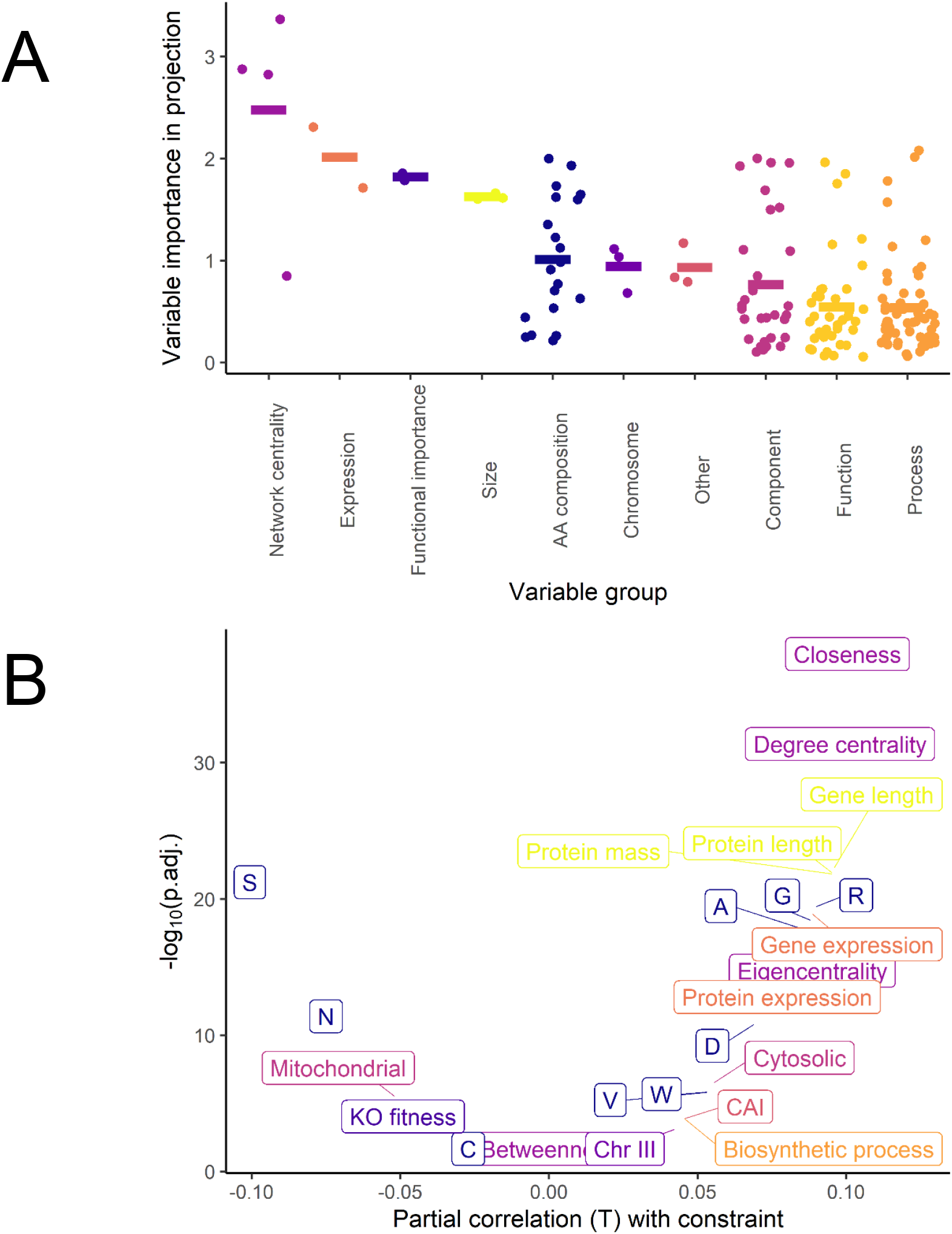
A) Variable importance in projection scores per variable group, showing that network centrality measures are the by far most influential variables in a partial least squares model of molecular evolution rates. Horizontal bars show group mean. B) Significant partial correlations with constraint, using Bonferroni correction to adjust for multiple comparisons. Colours refer to groups in A.

As previously discussed, one would intuitively expect that the importance of a gene for fitness would be the most important determinant of constraint. While this is not true in the fission yeast protein coding genome, and indeed not in most genomes studied (Zhang and Yang 2015), both essentiality (VIP = 1.8) and knockout fitness (VIP = 1.8) display relatively high VIP scores. However, of the two, only knockout fitness displays a significant, if weak (τ = -0.04), partial correlation with constraint. It has been suggested (Bergmiller et al. 2012) that the reason ‘essential’ genes can be lost in evolution is that other genes replace their function; perhaps such functional redundancy undermines the relationship between functional importance and rate of molecular evolution?

Sequence length – measured as gene or protein length – also correlates positively with conservation (τ = 0.09), with moderate VIP scores for each of gene length (1.7), protein length (1.6), and protein mass (1.6). This relationship has been reported in several organisms, including humans (Lipman et al. 2002) and *Saccharomyces cerevisiae* (Lipman et al. 2002; Bloom et al. 2006). Lipman and colleagues suggested that this is because of selective pressure to make proteins as short as they can be without disrupting function in order to reduce the cost of translation, meaning that both proteins with high functional importance and proteins with specific functional roles are likely to be longer. However, given that this variable remains significant even when functional role and importance have been adjusted for using partial correlation, this appears to be an insufficient explanation. It may be – to adapt the misfolding avoidance hypothesis (Drummond et al. 2005) – that longer genes are under greater selective pressure to avoid sequence changes as the risk of misfolding would be expected to increase with sequence length.

### Amino acid composition and functional role both show great heterogeneity in impact on constraint

Amino acid composition appears to have an influence on constraint that varies greatly depending on the amino acid. A few amino acids show higher VIP scores than functional importance, size, and even protein expression (fig. 1a). Some amino acids are correlated with higher constraint, others show a negative correlation, and most seem not to be significantly related to the rate of molecular evolution at all (fig. 1b). The changeabilities of amino acids are known to differ depending on the structural requirements of the protein domain – particularly, there is a considerable difference in amino acids composition between transmembrane and other protein domains (Tourasse and Li 2000). It is known that alanine and glycine, which relatively speaking strongly correlate with conservation, are highly enriched in Low Complexity Regions (LCRs) of proteins, which are known to be highly conserved (Ntountoumi et al. 2019). That said, some of the amino acids that show no correlation with constraint or even display negative correlation (e.g. serine), are also common in LCRs (Radó-Trilla and Albà 2012). Regardless, it appears likely that amino acid composition serves as a proxy for protein domains and regions that are under different selective pressures, in turn affecting constraint.

As is clear from figure 1, some gene ontology categories contribute strongly to increased or decreased constraints, whereas most categories appear to have a negligible influence. This may be because some GO categories might be less well defined, resulting in a bias in measured constraint towards well-defined categories, although one would expect a considerable heterogeneity for biological reasons as well. The only GO categories that correlate significantly with constraint are cytosolic expression, biosynthetic processes, small molecule metabolic activity, and mitochondrial expression. It is well known that intracellular proteins are more conserved (Liao et al. 2010), although the results here indicate that e.g. nuclear proteins would be less strongly correlated with constraint. Both biosynthesis and small molecule metabolic processes show an unsurprising positive correlation with constraint; this would be expected given their role in essential and evolutionarily conserved biological processes. Most of the GO slims with the highest VIP values (fig. 2) do not significantly correlate with constraint (fig. 1), suggesting that their high conservation is probably best explained by their correlation with other variables – such as gene expression. Most GO categories appear to fall far below the arbitrary VIP ‘significance’ cut-off of 1, suggesting that most have very little influence on gene conservation. It appears that a few functional role categories may have a causal relationship with sequence evolution constraint; in many cases, however, functional role is a proxy for other variables influencing constraint.

**Figure 2.**
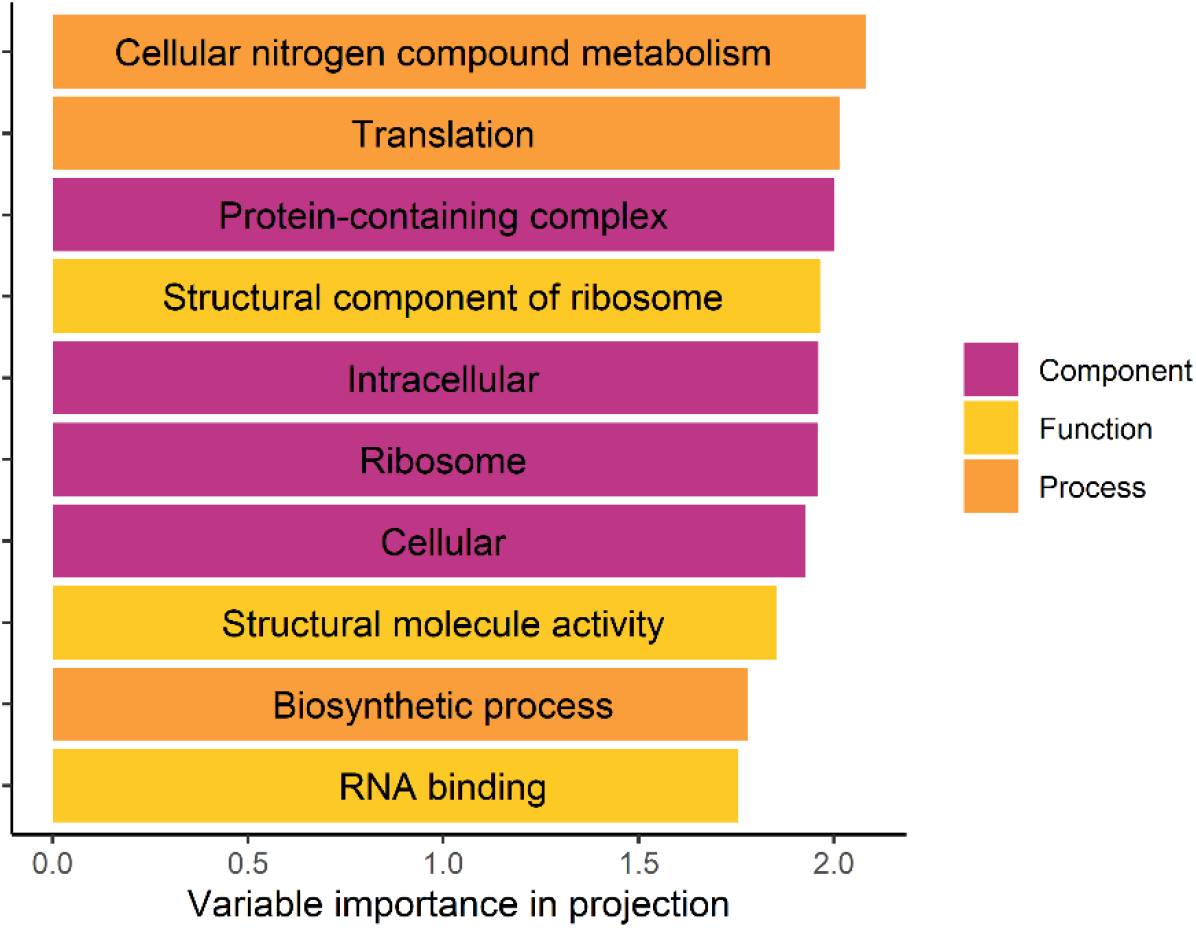
Then ten gene ontology slims with highest variable importance in projection (VIP) scores.

### Most of the variance in sequence evolution rate remains unexplained by these variables

Combined in a partial least squares model, these variables only explain 32.1% of the variance in constraint at the optimum number of model components. This shows that the model fails to adequately explain most of the variation in constraint. This could be due to a poor choice of modelling technique, but comparison with principal components regression (PCR), a highly similar method to PLS, shows that PLS explains more variance at far fewer components, with comparable root mean squared error (table 1). A random forest (Svetnik et al. 2003) regression model, a state-of-the-art machine learning technique suitable for high-dimensional noisy data, shows better performance (table 1) – but still fails to explain about 60% of the variance. Note that random forests are not necessarily suitable for robust and unbiased variable importance estimation (Strobl et al. 2007), which is why estimation of variable importance in the RF model has not been done here.

**Table 1.** A comparison of the performance of three regression techniques applied to sequence evolution constraint prediction.

| Regression model | RMSE <sup>1 2</sup> | Variance in constraint explained <sup>2</sup> | Number components |
| --- | --- | --- | --- |
| Partial least squares (PLS) | 0.167 | 32.1% | 3 |
| Principal components (PCR) | 0.169 | 30.7% | 74 |
| Random Forest (RF) | 0.153 | 42.9% |  |

<sup>1</sup> Root mean squared error. <sup>2</sup> Evaluation was performed on the 20% test set.

While the fit of the PLS model is not ideal, none of these three techniques manage to produce a model that explains even half of the variance in conservation. Given the high flexibility of random forest models in particular, it does appear unlikely that any commonly used regression technique would produce a model explaining much more than 50% variance. In other words, the high unexplained variance is a property of the data and is not just due to model choice. Notably, models of protein evolutionary rates in humans (Alvarez-Ponce et al. 2017) and *Arabidopsis* (Yang and Gaut 2011) have only been able to explain between 11% and 18% of variance in rates.

One feasible explanation for the low proportion of variance explained is poor data quality resulting from high technical noise – particularly of interactions and functional role data. Another possible reason is that some variables have not been explored, or not explored in sufficient depth. For instance, this study has analysed what the impact of which chromosome a gene is on has on constraint, ignoring finer details such as where on the chromosome – this data is available but difficult to include in modelling without overfitting – or location in the 3D genome. Similarly, individual GO terms rather than GO slims could be considered, but this would lead to difficulties in generalisable interpretation, as well as overfitting.

The more interesting explanation for the poor fit is that the rate of sequence evolution for a gene in the protein coding genome of fission yeast may be largely stochastic. This would be consistent with the neutral theory of molecular evolution (Kimura 1968), which states that most genetic variation is due to random drift. In this scenario rates of evolution should show considerable heterogeneity due to chance, resulting in large statistical noise. It appears likely that this would account for a considerable part of the missing variance. Determining how much of this is due to technical noise and missing data, how much is due to missing variables, and how much is due to stochastics may prove very difficult to determine. A systematic comparison across multiple species could provide useful insights about the variation in unexplained variance; this in turn might indicate the roles of data availability and quality, both of which vary greatly between organisms.

## CONCLUSION

Unlike what has been found in most organisms (Zhang and Yang 2015), it appears that network centrality is the variable that most strongly influences the rate of molecular evolution in fission yeast. Given that research from the past few years in the human genome (Alvarez-Ponce et al. 2017) has shown the high importance of network centrality as a variable reducing the rate of evolution, it is possible that this finding might be generally true across many organisms. Perhaps, until recently, this conclusion has been obscured by poor quality interactions data – whereas high quality gene expression data has been available for longer. Only a minority of variance in constraint is explained by all the variables combined, which is shown to be in part due to the limitations of the PLS regression technique, but largely due to the data itself. It consequently appears that the rate of molecular evolution in fission yeast may be largely influenced by variables not considered here, but also that stochastics may have a highly important role as well.

## DECLARATIONS

### Availability of data and material

All data are provided both in their original format and processed on Figshare (https://doi.org/10.6084/m9.figshare.c.5263523.v2). The dataset from Grech et al. was retrieved from Figshare (https://figshare.com/articles/dataset/gene-based_data/6265748). Protein data were retrieved from PomBase (ftp://ftp.pombase.org/pombe/Protein_data/PeptideStats.tsv and ftp://ftp.pombase.org/pombe/Protein_data/aa_composition.tsv). Protein interactions data were retrieved from STRING v11 (https://stringdb-static.org/download/protein.links.v11.0/4896.protein.links.v11.0.txt.gz).

### Code availabilitys

All R scripts are available both in the Figshare repository (https://doi.org/10.6084/m9.figshare.c.5263523.v2) and on GitHub (https://github.com/simonharnqvist/pombe)

### Ethics approval

Not applicable.

### Consent to participate

Not applicable.

### Consent for publication

Not applicable.

